# Mendelian Randomization Analysis of Circulating Adipokines and C-reactive Protein on Breast Cancer Risk

**DOI:** 10.1101/720110

**Authors:** T Robinson, RM Martin, J Yarmolinsky

**Author notes:** Corresponding author: Timothy Robinson, BMBS, PhD, MRCP, Bristol Cancer Institute, Horfield Rd, Bristol, United Kingdom, BS2 8ED.

## Abstract

Adipokines and C-reactive protein (CRP) have been proposed as molecular mediators linking adiposity to breast cancer (BCa). Mendelian randomization (MR) uses genetic variants as proxies for risk factors to strengthen causal inference in observational studies. We performed a MR analysis to evaluate the causal relevance of six circulating adipokines (adiponectin, hepatocyte growth factor, interleukin-6, leptin receptor, plasminogen activator inhibitor-1, resistin) and CRP in risk of overall and oestrogen receptor-stratified BCa in up to 122,977 cases and 105,974 controls. Genetic instruments were constructed from single-nucleotide polymorphisms robustly (*P*<5×10^−8^) associated with risk factors in genome-wide association studies. In MR analyses, there was evidence for a causal effect of hepatocyte growth factor on ER- BCa (OR per SD increase:1.17, 95% CI: 1.01-1.35; *P*=0.035) but little evidence for effects of other adipokines or CRP on overall or oestrogen receptor-stratified BCa. Collectively, these findings do not support an important etiological role of various adipokines or CRP in BCa risk.

---

Elevated body mass index (BMI) is an important modifiable risk factor for breast cancer (BCa)(1) and adipokines – cytokines and hormones released by adipose tissue- are potential molecular mediators linking excess adiposity to BCa(2–4). *In vitro* studies have demonstrated that two adipokines in particular – leptin and adiponectin – may have pro- and anti-proliferative effects on BCa cells, respectively, (5) and meta-analyses of observational studies support their opposing roles in BCa risk(6, 7). Observational studies have linked other adipokines including hepatocyte growth factor (HGF), interleukin-6 (IL-6), plasminogen activator inhibitor-1 (PAI-1), and resistin to BCa, albeit less consistently(8–10). Pre-diagnostic C-reactive protein (CRP), a systemic marker of inflammation that is partially synthesized by adipose tissue(11), has also been associated with BCa risk in prospective observational studies(12). Collectively, these findings suggest that pharmacological targeting of adipokines or CRP could be an effective strategy for BCa prevention among overweight and/or obese women. However, the causal nature of these risk factors in BCa risk is unclear as conventional observational analyses are susceptible to residual confounding and reverse causation, which undermine causal inference(13, 14).

Mendelian randomization (MR) uses genetic variants as instruments (“proxies”) for risk factors to generate more reliable evidence on the causal effects of these factors on disease outcomes(15, 16). The use of genetic variants as instruments minimises confounding and precludes reverse causation as germline genotype is largely independent of lifestyle and environmental factors and is fixed at conception. The power and precision of MR analysis can be increased by a “two-sample MR” framework in which summary genetic association data from independent samples representing genetic variant-exposure and genetic variant-outcome associations are synthesised in order to estimate causal effects(17).

Given uncertainty surrounding the role of various adipokines and CRP in BCa aetiology, we performed two-sample MR analyses to evaluate the potential causal role of circulating adiponectin, HGF, IL-6, leptin receptor, PAI-1, resistin, and CRP in overall and oestrogen receptor (ER)-stratified BCa risk.

Summary genome-wide association study (GWAS) statistics were obtained from analyses on 122,977 BCa cases (with further sub-analyses of 69,501 ER-positive (ER+) and 21,468 ER-negative (ER-) BCa cases) and 105,974 controls of European ancestry from the Breast Cancer Association Consortium (BCAC)(18). BCAC samples have the relevant ethical approval and genotyping was performed as previously described(18, 19).

Genetic instruments to proxy HGF, IL-6, leptin receptor, and resistin were constructed by obtaining individual *cis*-acting single-nucleotide polymorphisms (SNPs) robustly associated with these markers (*P*<5×10^−8^) in GWAS of individuals of European ancestry that were replicated in independent samples. *Cis*-variants (located ≤1MB of the transcription start site of the protein-coding gene) are more likely to have direct effects on protein levels than *trans*-variants (>1MB of the transcription start site of the protein-coding gene), minimising horizontal pleiotropy (an instrument influencing an outcome through one or more biological pathways independent to that of the exposure), a violation of the exclusion restriction criterion(16). For risk factors with ≥three independent (*r*^2^<0.01) *cis*- or *trans*-SNPs available as proxies (adiponectin, CRP, PAI-1), these SNPs were combined into multi-allelic instruments to increase the variance in the risk factor explained by the instrument. As sensitivity analyses for adiponectin, CRP, and PAI-1, causal estimates generated from multi-allelic instruments were compared with those obtained from instruments consisting of weakly correlated (*r*^2^<0.15) *cis-*variants to investigate horizontal pleiotropy in primary multi-allelic models.

*R*^2^ and F-statistics were calculated to examine the strength of our instruments, using previously reported methods(20). For instruments constructed using individual *cis*-variants, causal estimates were generated using the Wald ratio and standard errors were approximated using the delta method. For instruments constructed using ≥three independent variants, causal estimates were generated using inverse-variance weighted (IVW) random effects models to account for overdispersion in models (21). If underdispersion in a model was present, the residual standard error was set to 1. Sensitivity analyses for analyses employing multi-allelic instruments using weakly correlated *cis*-variants were performed using random-effects IVW models with adjustment for correlations between variants(22).

For each risk factor, the number of SNPs included in the instrument and estimates of instrument strength (*R*^2^ and F-statistics) are presented in **Table 1** with F-statistics ranging from 19.0-3872.7, suggesting that analyses were unlikely to suffer from weak instrument bias(23).

**Table 1.**
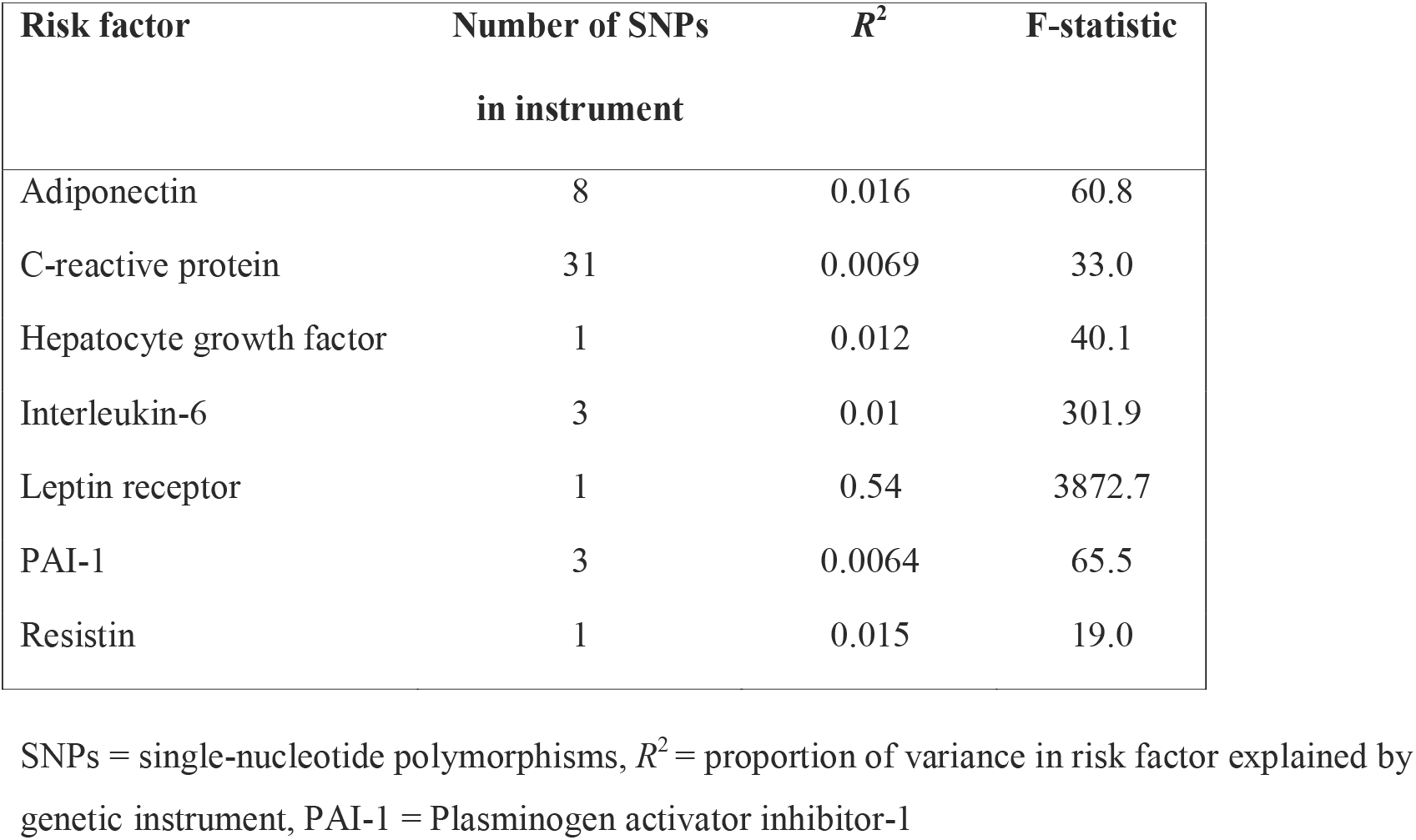
Number of SNPs included in instrument, estimate of the proportion of variance in risk factor explained by the instrument (*R*^2^), and F-statistic for each instrument, across all adipokines and C-reactive protein

In MR analyses, there was little evidence to suggest causal effects for any of the adipokines or CRP in overall breast cancer (**Table 2**). In ER status-stratified analyses, there was evidence for an effect of HGF on ER- BCa risk (OR per SD increase:1.17,95%CI:1.01-1.35;*P*=0.035). Findings for adiponectin, PAI-1, and CRP using *cis*-SNP instruments were consistent with those using multi-allelic instruments (**Supplementary Table 1**).

**Table 2.** Effect estimates per unit increase in adipokines or C-reactive protein on overall and oestrogen-receptor stratified breast cancer risk

| Risk factor | Overall breast cancer |  | ER+ breast cancer |  | ER- breast cancer |  |
| --- | --- | --- | --- | --- | --- | --- |
|  | OR (95% CI) | *P* value | OR (95% CI) | *P* value | OR (95% CI) | *P* value |
| Adiponectin | 1.06 (0.94-1.21) | 0.34 | 0.98 (0.84-1.14) | 0.80 | 1.19 (0.95-1.50) | 0.14 |
| C-reactive protein | 0.96 (0.88-1.05) | 0.38 | 0.96 (0.86-1.06) | 0.46 | 1.03 (0.87- 1.21) | 0.74 |
| Hepatocyte growth factor | 1.01 (0.93-1.10) | 0.77 | 1.01 (0.92-1.11) | 0.86 | 1.17 (1.01-1.35) | 0.035 |
| Interleukin-6 | 1.09 (0.96-1.25) | 0.18 | 1.12 (0.96-1.31) | 0.14 | 1.00 (0.79-1.27) | 0.99 |
| Leptin receptor | 1.00 (0.99-1.01) | 0.63 | 1.00 (0.99-1.01) | 0.81 | 1.00 (0.98-1.02) | 0.78 |
| PAI-1 | 0.97 (0.85-1.10) | 0.64 | 1.02 (0.87-1.19) | 0.81 | 0.95 (0.75-1.20) | 0.64 |
| Resistin | 1.04 (0.95-1.13) | 0.38 | 1.05 (0.95-1.17) | 0.33 | 1.01 (0.86-1.18) | 0.92 |

ER+ = Oestrogen receptor positive, ER- = Oestrogen receptor negative, PAI-1 = Plasminogen activator inhibitor-1, OR = Odds Ratio, 95% CI = 95%
Confidence Interval. Causal estimates represent the effect of a one unit increase in: natural log-transformed adiponectin, C-reactive protein, interleukin-6, and plasminogen activator inhibitor-1, and standardized hepatocyte growth factor, leptin receptor, and resistin

Contrary to some conventional observational studies (6–9, 12), our MR analyses using genetic variants as proxies found little evidence to support causal roles for various adipokines or CRP in BCa risk. Our data support a causal role of circulating HGF in ER- BCa risk that is consistent with *in vitro* studies suggesting a role of HGF in tumour cell proliferation, migration, and invasion (24, 25) and observational studies reporting a relationship of HGF levels with more advanced BCa staging and worse prognosis(10, 26, 27). While this result could be compatible with chance given the number of statistical tests performed, the alignment of findings from laboratory, observational, and genetic studies suggests the potential aetiological role of HGF in ER- BCa development.

Strengths of this analysis include the use of a two-sample MR framework that enabled increased statistical power and precision by exploiting summary genetic data from several large GWAS. There are several limitations to these analyses. First, since analyses were performed using summary genetic data in aggregate, this precluded stratification according to menopausal status. Second, though attempts were made to circumvent potential violations of MR assumptions in our analyses through the use of *cis*-acting variants as primary instruments and in sensitivity analyses, we cannot rule out the possibility that false negative findings may have arisen through horizontally pleiotropic pathways biasing our findings toward the null. Lastly, we were unable to examine possible non-linear effects of adipokines or CRP on BCa risk.

Overall, our findings suggest that several adipokines and CRP are unlikely to causally influence BCa risk. The potential aetiological role of HGF in ER- BCa warrants further investigation as a pharmacological target for BCa prevention.

