## Supplementary Table 1 for "Mendelian Randomization Analysis of Circulating Adipokines and C-reactive Protein on Breast Cancer Risk"

**Supplementary Table 1. Effect estimates per unit increase in adiponectin, C-reactive protein, and plasminogen activator inhibitor-1 on overall and oestrogen-receptor stratified breast cancer risk using conservative (*cis*-SNP) instruments***

| **Risk factor** | **Overall breast cancer** | **ER+ breast cancer** | **ER- breast cancer** |
| --- | --- | --- | --- |
|  | **OR (95% CI)** | **OR (95% CI)** | **OR (95% CI)** |
| Adiponectin | 0.97 (0.92-1.02) | 0.98 (0.92-1.05) | 0.98 (0.89-1.09) |
| C-reactive protein | 1.01 (0.95-1.07) | 1.02 (0.96-1.10) | 1.04 (0.94-1.16) |
| PAI-1 | 1.10 (0.93-1.30) | 1.16 (0.94-1.42) | 1.13 (0.83-1.53) |

SNP = single-nucleotide polymorphism, OR = Odds Ratio, 95% CI = 95% Confidence Interval, PAI-1 = Plasminogen activator inhibitor-1.

**Cis*-SNP instruments were constructed as follows: adiponectin was instrumented using four variants within *ADIPOQ* (rs6810075, rs16861209, rs17366568, rs3774261), C-reactive protein was instrumented using four variants within *CRP* (rs3093077, rs1205, rs1130864, rs1800947), and plasminogen activator inhibitor-1 was instrumented using one variant within *SERPINE* (rs2227631).

Causal estimates represent the effect of a one unit increase in: natural log-transformed adiponectin, C-reactive protein, and plasminogen activator inhibitor-1
